# Ancient DNA reveals that few GWAS loci have been strongly selected during recent human history

**DOI:** 10.1101/2021.04.13.439742

**Authors:** Corinne N. Simonti, Joseph Lachance

## Abstract

Genetic data from ancient humans has provided new evidence in the study of loci thought to be under historic selection, and thus is a powerful tool for identifying instances of selection that might be missed by methods that use present-day samples alone. Using a curated set of disease-associated variants from the NHGRI-EBI GWAS Catalog, we provide an analysis to identify disease-associated variants that bear signatures of selection over time. After accounting for the fact that not every ancient individual contributed equally to modern genomes, a Bayesian inference method was used to infer allele frequency trajectories over time and determine which disease-associated loci exhibit signatures of natural selection. Of the 2,709 variants analyzed in this study, 895 show at least a weak signature of selection (|s| > 0.001), including multiple variants that are introgressed from Neanderthals. However, only nine disease-associated variants show a signature of strong selection (|s| > 0.01). Additionally, we find that many risk-associated alleles have increased in frequency during the past 10,000 years. Overall, we find that disease-associated variants from GWAS are governed by nearly neutral evolution. Exceptions to this broad pattern include GWAS loci that protect against asthma and variants in MHC genes. Ancient samples allow us an unprecedented look at how our species has changed over time, and our results represent an important early step in using this new source of data to better understand the evolution of hereditary disease risks.

## Introduction

Many studies have been conducted to understand the genetic basis of diseases and other traits. Indeed, thousands of disease loci have been implicated in genome-wide association studies (GWAS) across human populations (Welter et al., 2014). Due to a balance between mutation and selection, alleles that are associated with Mendelian diseases tend to be rare (Weghorn et al., 2019). By contrast, alleles that are associated with complex diseases often segregate at intermediate frequencies (Timpson et al., 2018). GWAS loci are also enriched for derived, as opposed to ancestral, alleles that increase the risks of complex polygenic diseases (Gorlova et al., 2012; Lachance, 2010). There has been a great deal of interest in understanding how allele frequencies— and thus, traits—have changed over time; however, until recently, only modern genomes have been available. Analysis of modern human genomes has revealed that genes that confer resistance to infectious diseases have been targets of recent selection (Karlsson et al., 2014), and that natural selection contributes to differences in hereditary disease risks between human populations (Lachance et al., 2018; Marigorta et al., 2011). Classic debates about the relative importance of natural selection and genetic drift have also been rekindled (Jensen et al., 2019; Kern and Hahn, 2018). It has also been proposed that modern medicine has led to the relaxation of selection against mildly deleterious mutations and an increase in genetic load over recent human history (Lynch, 2016).

During the past decade, an increasing number of ancient genomes have been sequenced, some to very high read depth (Fu et al., 2014; Mathieson et al., 2015; Meyer et al., 2012; Narasimhan et al., 2019; Olalde et al., 2018; Prüfer et al., 2014). High quality genome sequences of our archaic hominin cousins have revealed previously unknown introgression events, and follow-up studies have shown how these introgressed regions have impacted modern human health and traits (Dannemann et al., 2016; Gunz et al., 2018; Simonti et al., 2016). Ancient DNA has also revealed that selection on the fatty acid desaturase genes *FADS1* and *FADS2*, which are relevant to lipid disorders, did not coincide with the Neolithic transition and the discovery of agriculture (Mathieson and Mathieson, 2018). Application of polygenic risk scores to ancient genomes has revealed that genetic disease risks have changed over the last 10,000 years (Berens et al., 2017). Furthermore, ancient human genomes have revealed that many variants associated with disease risk have increased in frequency during recent history (Aris-Brosou, 2019). Because ancient DNA enables times series data to be analyzed, it is now possible to determine the evolutionary histories of individual disease-associated loci. Large allele frequency changes during the last 10,000 years are indicative of disease loci that are under natural selection, while relatively flat allele frequency trajectories are indicative of disease loci that are evolving neutrally (Schraiber et al., 2016).

An open question in evolutionary medicine is whether the effects of genetic variation on disease risks are decoupled from fitness effects (Rodríguez et al., 2014). Here, we use 143 ancient human samples from 18 previously published studies to examine the role of selection on variants from the NHGRI-EBI GWAS Catalog. Most of the ancient samples analyzed here lived during the past 10,000 years. First, we correct for whether ancient genomes contribute to modern human populations. We then use time series data to infer allele frequency trajectories and selection coefficients for a curated set of 2,709 different disease-associated variants. We also examine whether protective alleles are more likely to be positively selected, and identify which diseases are enriched for signatures of weak selection. Collectively, these analyses reveal that most disease-associated loci are governed by nearly neutral evolution.

## Results

### Most GWAS variants do not show signatures of strong selection

For each disease-associated locus, we employed a Bayesian MCMC method (Schraiber et al., 2016) to infer allele frequency trajectories and selection coefficients that match time series data from ancient genomes. However, inference of natural selection requires an understanding of population continuity, or how much ancient individuals have contributed to modern genomes, if at all. As previous studies have found differences in contribution based on lifestyle (Haak et al., 2015), we grouped our ancient samples into one of three lifestyles: hunter-gatherers, agriculturalists, and pastoralists. We then calculated maximum possible genetic contribution of each of these groups to each variant using the method developed by Sjödin et al. (2014). Using this information, we could ensure we did not include ancient individuals who might skew allele frequencies in a way that did not reflect ancient demography (Figure S1).

Most GWAS loci have flat allele trajectories and selection coefficients that are close to zero. Of the 2,709 variants analyzed here, we find that 895 show at least a weak signature of selection (|*s*| > 0.001) and only 9 show a signature of strong selection (|*s*| > 0.01; Table 1). We also examined whether protective alleles were more likely to increase in frequency during recent history than risk-increasing alleles. A slight majority (52.6%) of putatively selected loci have protective alleles with positive selection coefficients. However, this trend was not statistically significant (471 of 895 variants, binomial test, *P* = 6.21 × 10^−2^). We also found that derived alleles are more likely to have positive selection coefficients than ancestral alleles at GWAS loci (528 of 895 variants; binomial test, *P* < 10^−7^). Taken together, our findings reveal that GWAS loci are largely governed by nearly neutral evolution.

**Table 1.** Selection coefficients inferred from ancient genomes.

| Variant type | Total number of variants | $\vert s\vert > 0.001$<br>(weak selection) | Protective allele selected | $\vert s\vert > 0.01$<br>(strong selection) | Protective allele selected |
| --- | --- | --- | --- | --- | --- |
| **All** | 2,709 | 895<br>(33.0%) | 471 of 895<br>(52.6%) | 9<br>(0.3%) | 4 of 9<br>(44.4%) |
| **MHC** | 148 | 76<br>(51.4%) | 36 of 76<br>(47.4%) | 4<br>(2.7%) | 0 of 4<br>(0%) |
| **Introgressed** | 24 | 8<br>(33.3%) | 4 of 8<br>(50.0%) | 0<br>(0.0%) | - |

Despite most variants appearing to evolve neutrally, exceptions exist for some disease-associated loci (Supplemental Table S1). For example, six of the nine variants under strong selection (s > 0.01) are associated with autoimmune diseases. One of them (rs13277113) is associated with systemic lupus erythematosus (Hom et al., 2008), and has been found by the GTEx Project (v8) to significantly affect expression of *BLK*, a gene involved in B cell development (Gauld and Cambier, 2004). Another of these variants (rs2270450) is associated with developing Hashimoto thyroiditis versus Graves’ disease (Oryoji et al., 2015). Curiously, this variant is significantly associated with expression of *TDRD6*, a gene involved in spermiogenesis as well as autoimmune polyendocrine syndrome (Akpınar et al., 2017; Bensing et al., 2007). The other four strongly selected variants associated with autoimmune diseases are found within the MHC region, which is known to be dense with rapidly evolving genes critical to immune function.

### Archaic introgression introduced some selected alleles at GWAS loci

Disentangling the effects of selection and archaic introgression can be difficult if one is analyzing modern genomes (Racimo et al., 2016). However, time series data from ancient genomes can sidestep many of these difficulties. Of the 2,709 disease-associated loci analyzed here, 24 contain introgressed alleles from Neanderthals. Of these 24 loci, nine introgressed variants have signatures of weak selection in the recent past (Figure 1). Some introgressed alleles appear to have reached moderate allele frequencies prior to being negatively selected, while other introgressed alleles appear to have only been positively selected during recent human history. Overall, we find that disease-associated alleles that have a Neanderthal origin have similar selection coefficients to other disease-associated alleles from the GWAS Catalog (Table 1). This is not wholly surprising, given that purging of strongly deleterious introgressed alleles occurred shortly after admixture (Petr et al., 2019), and introgressed alleles that are able to persist to the present day will have managed to avoid being filtered out due to Dobzhansky-Muller incompatibilities or other hybrid fitness effects.

**Figure 1.**
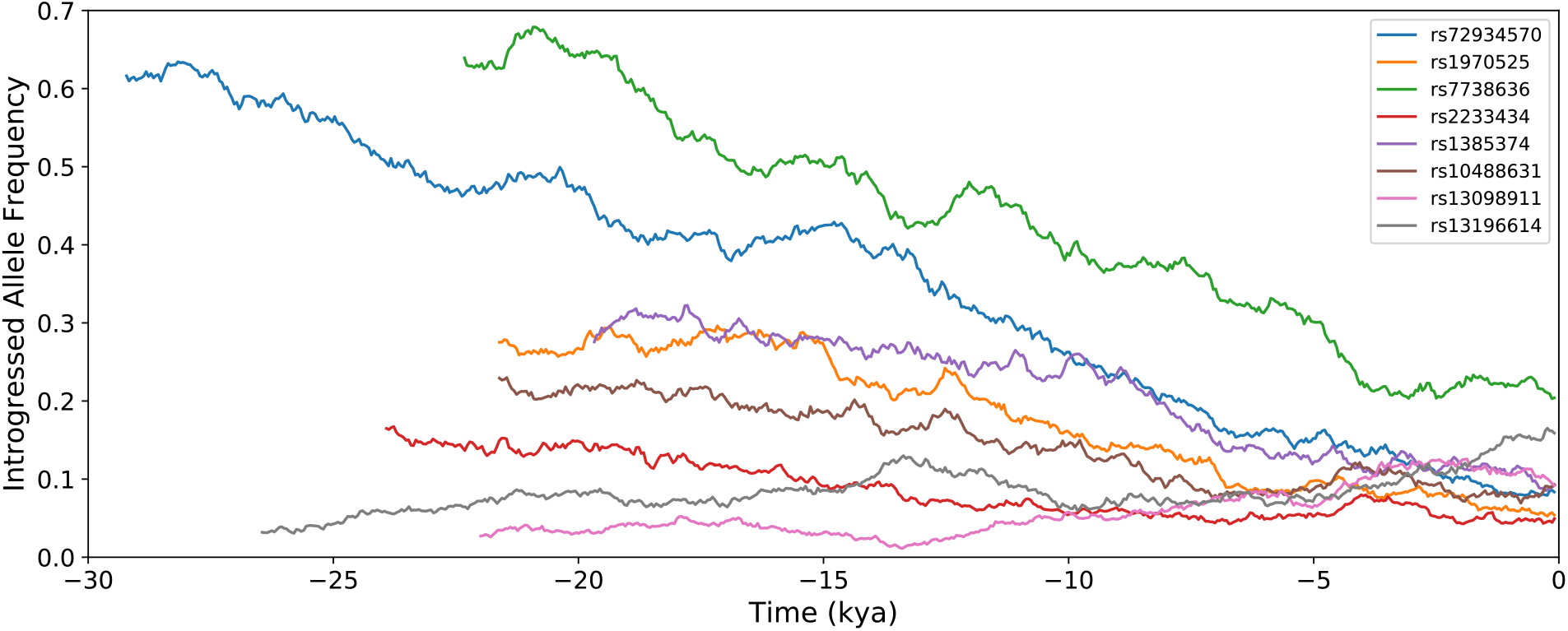
Some introgressed disease alleles are under weak selection. Inferred allele frequency trajectories of introgressed alleles with |*s*| > 0.001 are plotted here

### Most selected alleles at disease loci have small effect sizes

We examined whether selection coefficients are correlated with disease-related effect sizes. Given that our time series dataset exclusively contains samples from Europe and effect sizes are contingent on a host of factors including study population and environmental factors, this analysis focused on the subset of 1,910 variants that were identified in a GWAS that used samples of European ancestry (Figure 2). Overall, we find that there is no relationship between odds ratio (OR) and magnitude of *s* for non-MHC variants (*r* = −0.012). This pattern arises because many GWAS Catalog traits are largely irrelevant to fitness. For example, the variant with the highest OR analyzed here (rs3825942, OR=20.1) is associated with glaucoma, a disease which normally strikes long after reproductive age. Although the variants in Figure 2 are associated with a heterogenous set of diseases, our results suggest that the selection coefficients are largely decoupled from the effects of GWAS loci on disease risks.

**Figure 2.**
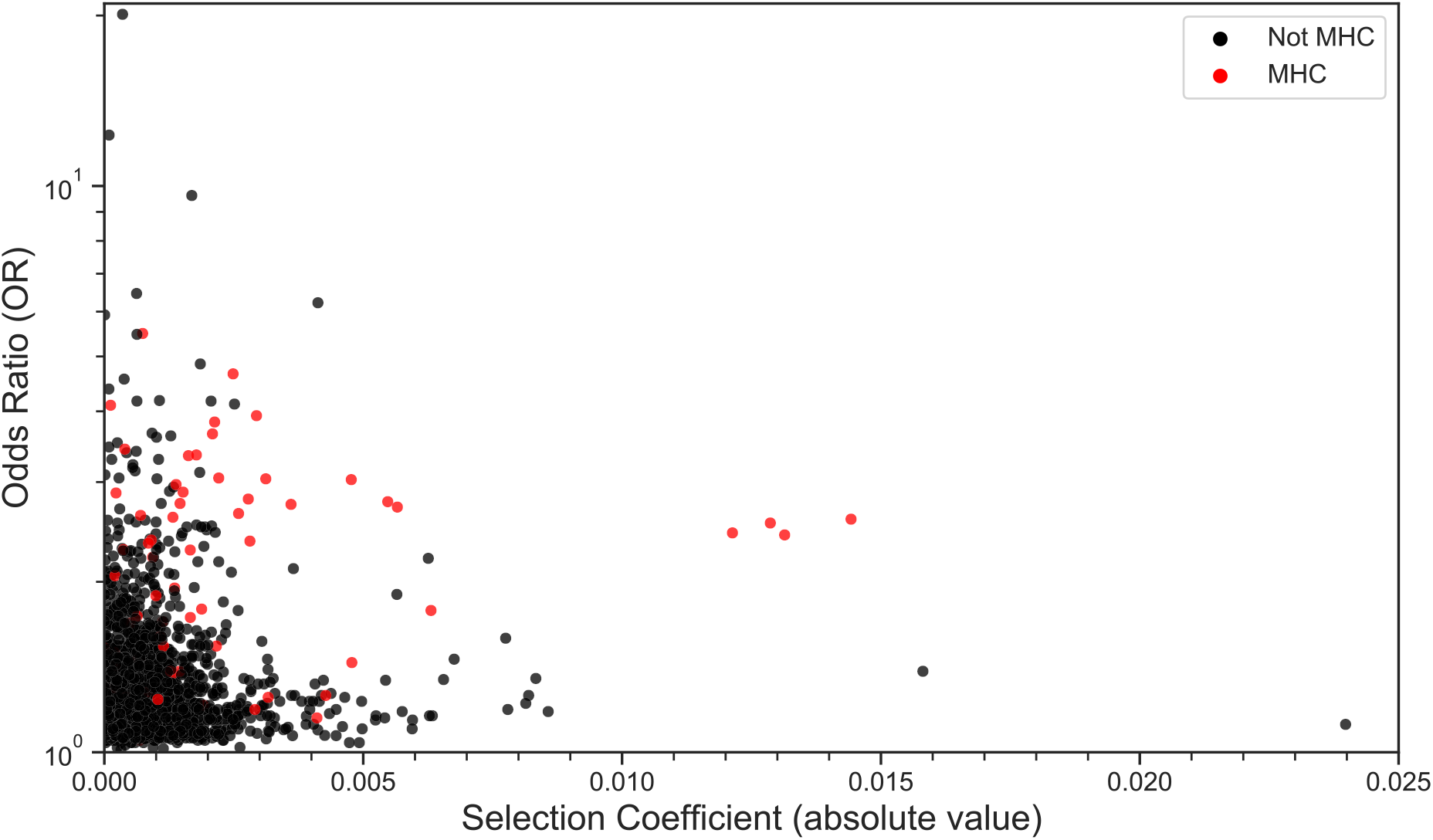
GWAS effect sizes are generally unrelated to selection coefficients. We plotted the odds ratios (OR) of GWAS loci verses inferred selection coefficients for 1,910 disease-associated variants that were identified in a GWAS that used samples of European ancestry (MHC variants: red, non-MHC variants: black).

### Variants in the MHC region show stronger signatures of selection

Compared to other disease-associated alleles, MHC variants show a strikingly different pattern. Disease-associated variants in the MHC region on chromosome 6 have larger effects sizes than disease-associated variants that are not in the MHC region (mean MHC OR: 2.11, mean non-MHC OR: 1.36, t test *P* = 4.12 × 10^−19^). Disease-associated variants in MHC genes also show a modest positive correlation between *s* and OR (*r* = 0.167). Variants located in the MHC region were significantly more likely to show signatures of weak selection than GWAS variants found in other parts of the human genome (Table 1, hypergeometric *P* < 10^−6^). Disease-associated variants in the MHC region were also significantly depleted of selection for protective alleles (hypergeometric *P* < 10^−6^). Given the highly pleiotropic nature of the immune system, it is possible that alleles that increase risk for some immune diseases have been historically protective against others unrepresented in the GWAS Catalog (i.e., infectious diseases).

### Disease-specific tests for enrichment of selection signatures

Although we caution against overinterpreting disease-specific results (due to genetic hitchhiking and the pleiotropic nature of variants analyzed in this paper), multiple diseases were enriched for signatures of weak selection. Idiopathic membranous nephropathy, an autoimmune disease that severely impacts kidney function, showed the strongest signature of selection, with 17 of 19 variants under at least weak selection (Figure 3A; hypergeometric *P* < 1.00 × 10^−6^). Unsurprising given the strong immune component to this disease, all but one of the variants associated with this disease are found in the MHC region, which likely contributes to enrichment for selected alleles that are associated with idiopathic membranous nephropathy. Curiously, alleles that protect against this disease have tended to decrease in frequency during recent human history (Figure 3B; hypergeometric *P* = 4.76 × 10^−4^). As stated above, this result could represent either genetic hitchhiking or pleiotropic effects. The next strongest enrichment signal was from asthma, with 26 of the 49 variants associating with it being under at least weak selection (Figure 3A; hypergeometric *P* = 2.12 × 10^−3^). We found that alleles that protect against asthma were strongly enriched for being under positive selection (Figure 3B; 20 of 26 selected alleles were protective; hypergeometric *P* = 8.94 × 10^−3^). This result was robust to the exclusion of MHC variants. A third disease that showed enrichment for signatures of weak selection was acne, with 7 of its 10 associated variants under selection (hypergeometric *P* = 1.79 × 10^−2^; Supplemental Table S2).

**Figure 3.**
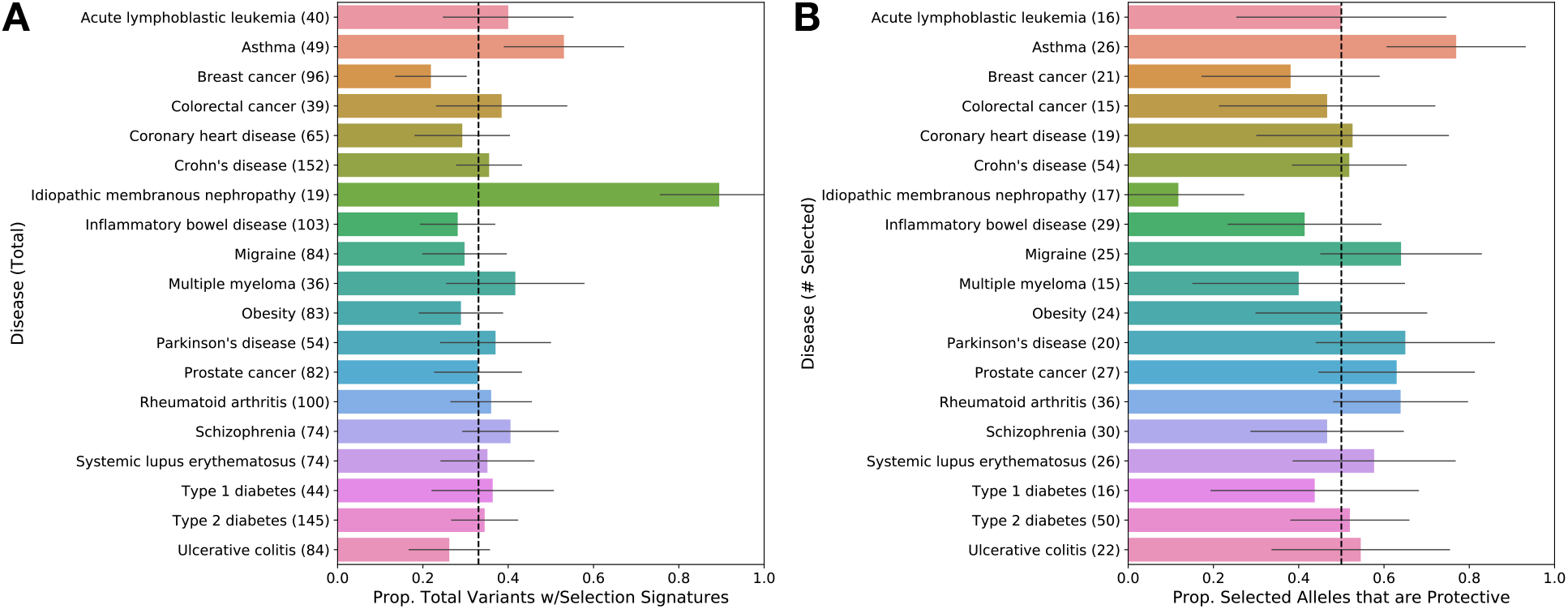
Few diseases are enriched for signatures of selection. Dashed vertical lines indicate mean values for all disease-associated alleles analyzed in this study. **(A)** For diseases with at least 15 variants under weak selection, the proportion of variants with |*s*| > 0.001 is plotted. Details about other diseases can be found in Supplemental Table S2. **(B)** The proportion of selected variants where the protective allele has a positive selection coefficient.

In addition, we found three diseases that were significantly depleted for signatures of weak selection (Supplemental Table S2). The strongest signal was for breast cancer (Figure 3B; 21 of 96 variants; hypergeometric *P* = 1.20 × 10^−2^), followed by major depressive disorder (10 of 55 associated variants; hypergeometric *P* = 1.27 × 10^−2^), and addiction (3 of 23 variants; hypergeometric *P* = 2.96 × 10^−2^). Taken together, these results reveal that not all diseases are equally affected by selection, even when selection does not appear to be broadly acting.

Given that severity and when a disease manifests determine its fitness impact, we also examined whether diseases with an early age of onset have different characteristics than diseases with a late age of onset. Overall, 55.9% of alleles that are associated with increased disease risks in our dataset were derived, as opposed to ancestral (binomial *P* < 1.00 × 10^−6^). This trend was more pronounced in diseases whose onset was before the age of 18 (hypergeometric *P* = 3.50 × 10^−5^; Figure 4A). Despite this effect, age of onset does not seem to impact the likelihood of a variant being under selection (Figure 4B). However, there is a slight bias in favor of the protective allele for both the 0-18 age of onset (binomial *P* = 0.047) and the 19-40 age of onset disease variants (binomial *P* = 0.042; Figure 4C).

**Figure 4.**
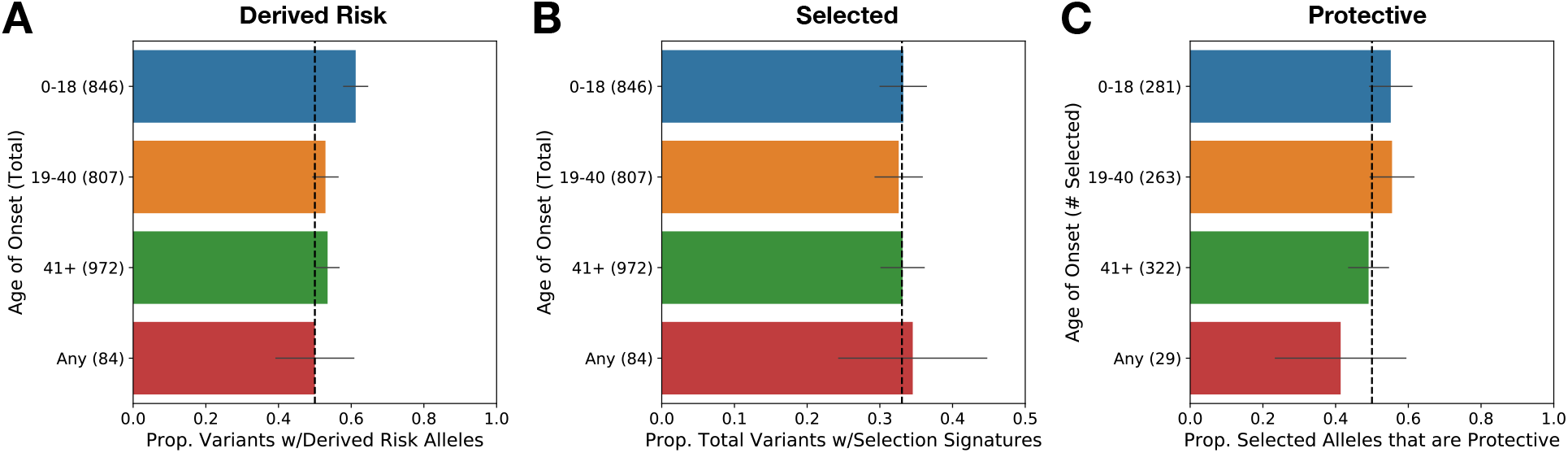
Comparisons between GWAS loci that are associated with diseases that have different ages of onset. **(A)** Proportion of variants where the risk-increasing allele is the derived allele. The dashed line represents the null expectation (50%). **(B)** Proportion of variants under at least weak selection (|*s*| > 0.001). The dashed line represents the overall variants under weak selection (33.0%). **(C)** Proportion of selected variants where protective alleles have increased in frequency over recent human history. The dashed line represents the null expectation (50%).

## Discussion

We find that the majority of GWAS variants do not appear to be under strong selection, and that selection coefficients (*s*) and odds ratios (OR) are uncorrelated. Allele frequency changes at many disease-associated loci may be due to the indirect effects of genetic hitchhiking, i.e., selection acting at closely linked loci. Because of this, our results suggest an “upper bound” for the amount of selection experienced by GWAS variants. In general, we find that our results reinforce the notion that diseases do not necessarily impact fitness. Our study also demonstrates the utility of time series data to infer the evolutionary histories of individual disease-associated loci.

Disease phenotypes are obvious targets for natural selection, so the general lack of variants under selection in this set of GWAS loci at first seems unexpected. However, a previous study estimating selection coefficients from C-scores showed a remarkably similar proportion of GWAS variants under weak selection (Racimo and Schraiber, 2014). Indeed, there are many reasons why we would not expect many of these variants to be under strong directional selection in the timescale examined. First, many of these diseases are unlikely to impact fitness, a scenario supported by our result that OR and the magnitude of *s* are uncorrelated for most variants. Of the 178 diseases considered here, 20 are late-onset (age of onset of 60 or later) and while they shorten lifespans, they do so well after reproductive age. Barring strong effects from scenarios such as the grandmother hypothesis (Williams, 1957) and presuming these variants substantially affect no other fitness-impacting traits, this makes this group of disease variants essentially invisible to selection. The fact that these diseases show no marked depletion in the proportion of variants under weak selection when compared to diseases with earlier ages of onset could imply that genetic hitchhiking or pleiotropy may be obscuring expected patterns of selection.

Another possibility is that selection has acted on disease traits, but the polygenic nature of the disease means individual loci affecting risk might be close to neutrality. Indeed, this is expected as many GWAS variants are segregating at allele frequencies that no strongly deleterious variants could ever reach. Previous studies have demonstrated negative correlations between allele frequency and effect size, even when study power was taken into account (Park et al., 2011). This suggests that purifying selection constrains evolutionarily deleterious alleles, supporting the idea that variants in the GWAS Catalog have not historically had large fitness effects. Another possibility is that due to environmental changes, some of these diseases have become more prevalent in recent history (so-called diseases of modernity). This would cause selection to only be able to act on them recently, outside the timescale of our analyses. These scenarios do not represent an exhaustive list of possibilities, nor are they mutually exclusive.

One exception to this overall pattern is that many putatively selected variants are found in the highly polymorphic MHC region of the genome. Genes in this region impact resistance to infectious disease, mate choice, kin recognition, reproductive success, among others (Bernatchez and Landry, 2003; Piertney and Oliver, 2006). As such, they have been frequent targets of recurring balancing and positive selection. When considered, these theories make it clear why GWAS variants overall would be depleted of signatures of selection, apart from MHC variants. It is worth noting the highly repetitive and highly polymorphic nature of this region makes genotyping in the MHC region notoriously difficult, so it is possible that this has introduced bias in our results (Kiyotani et al., 2017). However, there is no a priori reason why difficulty genotyping would consistently result in an excess of signal for selection.

Interestingly, diseases whose onset is during pre-reproductive or reproductive age are slightly more likely to have protective alleles that are positively selected. A prime example of this involves variants affecting asthma, which are strongly enriched for selection overall, and positive selection on protective alleles in particular. Asthma is a chronic inflammatory disorder of the airways primarily affecting children whose incidence has increased substantially over recent decades (Akinbami et al., 2015), which has been partially attributed to both the hygiene hypothesis as well as increasing air pollution through urbanization (Bowatte et al., 2015; WHO, 2019). These factors together suggest that the environmental risk for asthma has been on the rise over recent human history, with commensurate increases in selective pressure resulting in the strong enrichment for positive selection on protective alleles that we observed.

It is worth mentioning that the approach taken here is subject to the limitations of the GWAS data to which it has been applied. While incredibly useful, GWAS are not easily conducted on certain diseases that likely carry great importance to fitness, such as infectious diseases or diseases and health events that result in patient death before they can be enrolled in a study (i.e., myocardial infarction, stroke). Our approach here is well suited to examining the common variants typically found in GWAS, but it is ill-suited to rare or family-specific variation. Additionally, the timescales we can examine using this approach are defined by the density and timeline of ancient samples. As many of our samples are from the last 10,000 years and the youngest is from ∼1,208 years ago, very recent allele frequency shifts are difficult to detect. It is possible that some variants may have been under episodic selection, but our method is unable to detect selection if allele frequency trajectories are U-shaped.

## Conclusion

Our study shows the utility of ancient genomic data and time series analyses in understanding the effects of selection versus neutral processes on complex disease risk. These results contribute to the growing body of data on how selection has shaped the human genome, as well as the importance of disambiguating fitness effects from disease status. It is also worth noting that incorporating information from ancient samples can help give context in situations where traditional markers of selection (linkage disequilibrium, singleton density, etc.) are obscured due to the history of the region. This is a particular source of trouble in regions such as haplotypes introgressed from ancient hominins, whose history violates many assumptions of methods using modern data alone (Vernot and Akey, 2014). As more high-quality ancient genomes are collected from around the world, our understanding of evolutionary medicine will continue to grow.

## Materials and Methods

### GWAS variant and ancient DNA curation

Data was downloaded and curated as described in (Berens et al., 2017). Briefly, variants were downloaded from the GWAS catalog (accessed 28 September 2016) and associations with benign phenotypes removed. Specific variant associations were removed from consideration if two variants associated the same disease and were within 100 kb of each other, with a preference for maintaining the variant-phenotype association with the lower p-value. The risk allele for each variant was classified as derived or ancestral based on data from the 1000 Genomes Project (Auton et al., 2015). We used genomic data from 143 ancient individuals from 18 studies (see (Berens et al., 2017) for further details on the curation of this data). As time series data requires a continuous population, non-European ancient human samples (e.g., Mota and Anzick) and archaic hominin Neanderthal and Denisovan samples were not included from the larger set described in Berens et al (2017).

### Age of onset determination and phenotype collapsing

When available, we used the disease information collated by Google from multiple data sources (for more information, see: https://support.google.com/websearch/answer/2364942). Onset was broken down into seven age ranges: 0-2, 3-5, 6-13, 14-18, 19-40, 41-60, and 60+. For each disease, we used the age range determined by the highest frequency of ages affected. When multiple age ranges were equally likely, we used the youngest age range. For the diseases that Google did not have this information, we used either age of onset data or the age of study participants from the studies contributing to the GWAS Catalog (Welter et al., 2014). For phenotypes that lacked an age of onset (allergic reactions, injuries, etc.), we used the category “Any.” In addition, we collapsed similar phenotypes into an umbrella phenotype (Supplemental Table S2). For example, all symptoms associated with type II diabetes were collapsed into the type II diabetes umbrella category.

### Population continuity and contributions to modern genomes

We used a method developed by Sjödin et al (2014). Ancient individuals were assigned into three groups based on lifestyle: agriculturalists, hunter-gatherers, and pastoralists. For each variant, we used this program to determine the maximum possible genetic contribution of individuals from each lifestyle to modern EUR individuals. Maximum contribution was calculated by decile using a uniform prior and scaled admixture time of 0.01 (5000 years ago, given a generation time of 25 years and N_e_ of 10,000). This method was shown to be robust to a difference in the time of admixture compared to the age of the sample as well as variation in sample age. Consistent with previous findings (Sjödin et al., 2014), similar results were observed when using the Griffiths distribution, which incorporates ancestral and derived state information.

### Allele frequency trajectory and selection inference

We used a method developed by Schraiber et al (2016). For variants where the maximum contribution of individuals of a particular lifestyle was less than 100%, we down-sampled the individuals from that lifestyle by enforcing a chance of inclusion equal to the maximum contribution of that lifestyle. Individuals were then binned by age (>9500, 9500-8000, 8000-6500, 6500-5000, 5000-3500, and <3500 years ago). Each time point was determined by calculating the median age of all individuals included in the bin. Time was then scaled by generation time (25 years) and N_0_ (10,000). The program was then run twice for each variant, once using a demographic model of constant population size, and the other including the out of Africa bottleneck and subsequent exponential growth - parameters used from Gravel et al (2011). Growth was modelled as a step-wise exponential increase in population size every 2500 years until a population size of 20,000 was reached, after which population size was held constant through to present day. Variants were only included for analysis if at least 20 ancient individuals total contributed over at least two time points (determined after individuals were down-sampled). In instances of individual down-sampling, the same individuals were used for each variant for both constant and exponential runs. The final selection coefficient (*s*) was determined by averaging the *s*_*1*_ (selection on heterozygote) and 1/2 *s*_*2*_ (selection on homozygote) values generated by this method. Due to overestimation of *s* when the derived allele frequency started very high in ancient populations, we repolarized the test to treat an ancestral allele as if it was derived when the derived allele frequency at the first time point was greater than or equal to 75%.

### Introgressed and MHC variant identification

We determined the introgression status of variants using the haplotypes and variants identified in a study of Neanderthal and Denisovan introgression across modern Eurasian and Melanesian groups (Vernot et al., 2016). Coordinates defined by the Genome Reference Consortium were used to specify the MHC region (hg38 chr6:28,510,020-33,480,577). When necessary, we used liftOver to convert data from hg38 to hg19 or vice versa (Hinrichs et al., 2006).

### Binomial and hypergeometric enrichment

We used the numpy module in python to generate binomial and hypergeometric distributions. In all cases, we simulated 1,000,000 draws. Unless otherwise stated in the text, hypergeometric distributions were generated based on the results from the overall set of 2,709 GWAS variants with selection information.

## Acknowledgements

We thank Josh Schraiber and members of the Lachance Lab for their helpful feedback on this manuscript. This work was supported by NIH grant R35GM133727.

