## Supplementary material for "Ancient DNA reveals that few GWAS loci have been strongly selected during recent human history": Figure S1

### Supplemental Figure S1

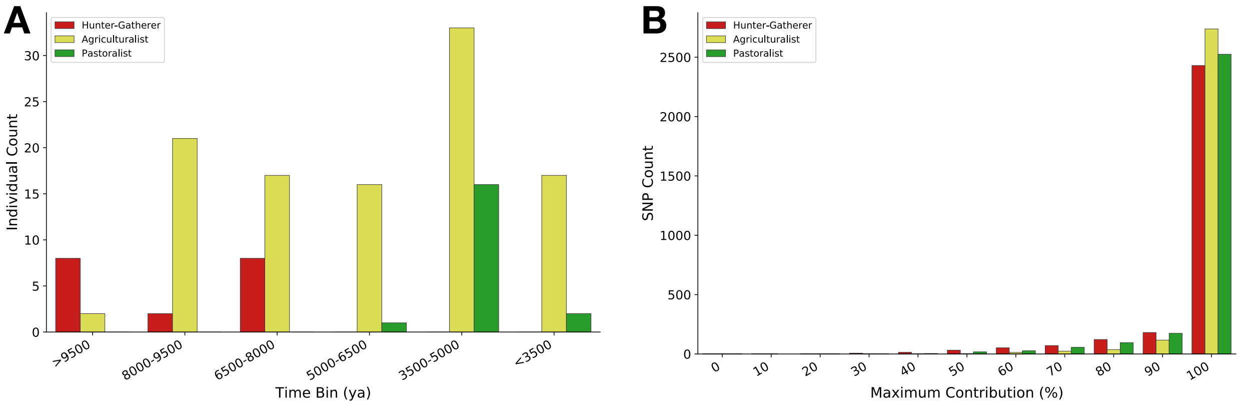


**Figure S1**. **Subsistence patterns impact ancient contributions to modern populations. (A)** The distribution of ancient samples across time bins. **(B)** Maximum contribution of ancient samples (grouped by lifestyle) to modern Europeans for each disease-associated locus in our study. This information was used to down-sample ancient individuals when inferring allele frequency trajectories and selection coefficients. Ancient agriculturalists contribute more to modern European genomes than ancient hunter-gatherers or pastoralists.
